# Flexibility Drives Information Flow in Proteins: Fluctuation Potential Gradients Dictate Directional Entropy Transfer

**DOI:** 10.64898/2026.08.14.744694

**Authors:** Fatma Senguler Ciftci, Burak Erman

## Abstract

Allosteric communication in biomacromolecules is fundamentally governed by thermal fluctuation gradients, yet standard Gaussian Network Models (GNMs) treat atomic contacts as uniform, binary couplings without differentiating core constraints from solvent-exposed surface flexibility. Here, we present an analytical matrix framework that incorporates continuous distance-dependent weighting into the Kirchhoff matrix *L*. This formulation captures the steep steric constraints of hydrophobic core packing versus peripheral surface loops while strictly recovering the classic unweighted GNM as a high-temperature limit (*T* →∞). Using Schur complements of partitioned joint covariance matrices, we show that conditional fluctuation variances and higher-order entropy-transfer terms reduce analytically to exact ratios of submatrix determinants (covariance minors), eliminating the need for fitting parameters or molecular dynamics trajectories. Applied to KRAS (PDB: 6GOD), this framework constructs an integrated directional entropy-transfer asymmetry map. Order-1 minors (*h*(*i*) = *K*_*ii*_) establish a single-node fluctuation potential gradient, while order-2 minors (*R*_*ij*_) define pairwise channel bandwidths. Higher-order minors show multi-body spatial coupling: order-4 minors identify rigid core residues such as Phe156 as strategic interlobe relay hubs linking Lobe 1 and Lobe 2, and an order-3 triad cooperation index demonstrates that signal transmission from Switch II (Gln61) to Gly60 and Phe156 converges on a single, mechanically integrated allosteric sector. By deriving directional information flow directly from experimental atomic displacement parameters, this approach establishes a rigorous, computationally efficient framework for mapping allosteric networks across structural ensembles.

## 1. Introduction

Proteins undergo continuous thermal fluctuations about their mean equilibrium conformations. In statistical thermodynamics, these thermal motions are not independent isotropic noise, but collective spatial fluctuations whose covariance structure is dictated by the underlying network topology. Recent breakthroughs in diffuse X-ray scattering and structural ensemble analysis, most notably pioneered by Meisburger et al. [1] and Xu et al. [2], have provided direct experimental measurement of these continuous displacement correlations, proving that macromolecular dynamics are governed by extended, cooperative fluctuation fields. The Gaussian Network Model formalizes these physical observations by representing the protein as a graph whose weighted Kirchhoff (graph Laplacian) matrix, *L*, generates the theoretical equilibrium covariance matrix *K* = *L*^+^, where *L*^+^ denotes the Moore– Penrose pseudoinverse of *L*. This covariance operator carries deep statistical-mechanical structure. It serves simultaneously as the Green function of the network, the covariance matrix of the fluctuation field, and a positive-semidefinite Gram matrix that embeds every residue as a vector in an abstract fluctuation Hilbert space equipped with effective distances, *R*_*ij*_. In a recent investigation, the present authors showed that the algebraic minor hierarchy of *K* generates a complete sequence of residue-subset invariants, effective distances *R*_*ij*_, cooperation indices, and multi-body volume measures that define allosteric communication across increasing levels of structural organization [3].

While the static covariance matrix *K* quantifies spatial correlations, correlation alone does not establish directionality or information transport. Two residues may exhibit strong equilibrium correlations without one exerting a directed influence on the future dynamical state of the other. To capture directional dependency, Schreiber introduced transfer entropy as a model-free, information-theoretic measure of directed information flow between stationary processes [4]. Subsequent developments by Hacisuleyman and Erman [5], alongside the formal equivalence proven by Barnett, Barrett, and Seth [6] for continuous Gaussian systems, established that transfer entropy in linear Gaussian networks can be evaluated analytically from the underlying network topology, bypassing the extensive sampling required by molecular dynamics trajectories [7, 8].

However, a fundamental physical question remains: what is the precise origin of directionality in an equilibrium fluctuation process? While recent computational investigations into domain allostery highlight the role of structural heterogeneity and distinct conformational sub-states [9], an open question is whether directional signal propagation requires distinct structural transitions or arises intrinsically from linear thermal fluctuations. Strictly speaking, equilibrium microscopic reversibility forbids a net probability current: every fluctuation trajectory is statistically balanced by its time-reversed counterpart. Directionality can nevertheless appear at the level of structural response or information flow because of anisotropic energy landscapes, unequal relaxation times, kinetic barriers, and coarse-grained observables. Genuine directed transport or cyclic flux, however, requires broken detailed balance and therefore an external source of free energy or another mechanism of nonequilibrium symmetry breaking. In accordance with Callen’s formulation of the fluctuation-dissipation theorem [10], an equilibrium system possesses no external thermodynamic driving force or non-equilibrium flux; its relaxation dynamics are governed entirely by the spatial distribution of equilibrium variance. The directional asymmetry of transfer entropy in a linear network is therefore not a thermodynamic heat dissipation or a conformational transition, but an intrinsic asymmetry in conditional predictability. Knowledge of the present state of an upstream residue reduces uncertainty about the future state of a downstream residue by a different amount than the reverse, precisely because the spatial distribution of dynamical memory is heterogeneous across the network.

In the present work, we show that this analytical conditional entropy framework simplifies, without structural approximation, to an intrinsic discrete potential gradient. We prove that the diagonal elements of the pseudoinverse covariance matrix, *h*(*i*) = *K*_*ii*_, define a single-valued residue fluctuation potential whose spatial differences ∇*h*(*i, j*) = *h*(*j*) − *h*(*i*) dictate the net directional flow of information in the short-time regime. Remarkably, the degree terms of the Laplacian cancel identically in the directional asymmetry, leaving a purely potential-driven field.

Since harmonic spatial variance maps directly onto crystallographic B-factors via classical equipartition, the theoretical node potential *h*(*i*) provides an empirical baseline for evaluating directional fluctuation gradients directly from experimental atomic displacement parameters. Without fitting empirical force-field parameters, specifying explicit relaxation time-lags, or running molecular dynamics trajectories, the directional source-and-sink topology of a protein can be read directly from deposited structures in the Protein Data Bank [11].

To interpret these directional maps, we establish *h*(*i*) as the foundational first-order level of the Laplacian minor hierarchy. While *h*(*i*) governs the driving potential for pairwise asymmetry Δ*T*_*ij*_, higher-order minors, such as the order-two effective distance 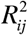 and order-three cooperation index χ_*i*; *jk*_, dictate channel bandwidth, multi-target pathway overlap, and structural relay hubs. The Laplacian minor hierarchy thus provides a unified structural language for decoding complex patterns in empirical asymmetry maps. We demonstrate this framework on oncogenic KRAS (PDB: 6GOD) [12]showing that theoretical predictions closely track experimental B-factor gradients and align with recent combinatorial Hodge decompositions that confirm the irrotational, potential-driven nature of protein information flow [13].

## 2. Theory

### 2.1 Structure-Induced Fluctuation Field and the Node Potential

In accordance with the statistical mechanics of Gaussian network models [14, 15], thermal fluctuations about equilibrium are governed by the contact topology of the residue interaction graph. The weighted Kirchhoff matrix, *L*, equivalent to the graph Laplacian, has elements defined by:

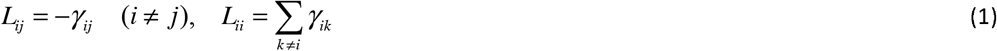

where the pairwise interaction weight *γ*_*ij*_ depends on the spatial distance *d*_*ij*_ between the *C*_*α*_ atoms of residues *i* and *j* :

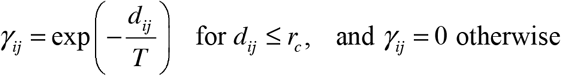

As discussed in our recent work [16], this continuous distance-dependent weighting differentiates short-range core interactions from peripheral surface contacts, a distinction absent in standard topological networks, while recovering the unweighted Gaussian Network Model (GNM) as a high-temperature limit (*T* →∞).

The Kirchhoff Laplacian matrix *L* is real, symmetric, and positive-semidefinite, possessing a single zero eigenvalue corresponding to rigid-body translational invariance. The equilibrium covariance matrix of residue fluctuations, *K*, is given by the pseudoinverse of *L* :

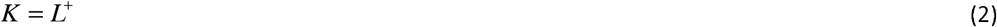

By formulation of the fluctuation-dissipation theorem and statistical mechanics of Gaussian chains [10, 17], *K* completely determines the spatial correlations and local variances of the network. Because *K* is positive-semidefinite, it defines a Gram matrix equipped with an inner product, *K*_*ij*_ = ⟨**v**_*i*_, **v**_*j*_ ⟩, embedding every residue *i* as a vector **v**_*i*_ in an abstract fluctuation Hilbert space [18]. The construction of this Hilbert space, and the proof that *R*_*ij*_ is a valid squared distance within it, are given in Appendix C.

The diagonal elements of *K* define a scalar residue-specific fluctuation potential, *h*(*i*) :

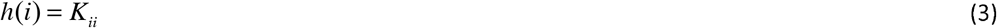

Physically, *h*(*i*) is proportional to the mean-square displacement of residue *i* and is directly proportional to its crystallographic B-factor. Residues in tightly constrained, densely packed environments exhibit small values of *h*(*i*), whereas flexible residues in weakly constrained loops exhibit large values of *h*(*i*). We regard *h*(*i*) as the local dynamical potential governing the conformational entropy distribution of the network.

The intrinsic geometry of this fluctuation space is characterized by the effective distance, *R*_*ij*_, which quantifies the relative mobility between residue pair (*i, j*) :

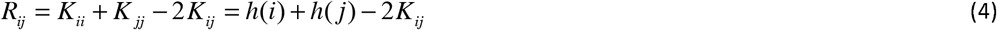

The effective distance *R*_*ij*_ serves as the natural metric of network flexibility, replacing physical Euclidean distance with topological resistance distance.

### 2.2 Langevin Dynamics and Representation Dualities

The time evolution of network fluctuations **Δr**(*t*) is modeled by the overdamped Langevin equation:

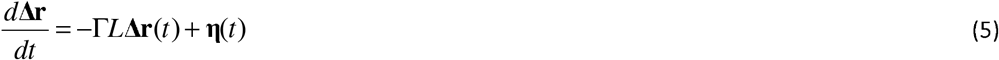

where Γ is a kinetic friction constant and **η**(*t*) represents Gaussian white noise satisfying

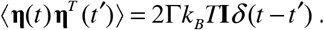

To account for global translational zero-modes, fluctuation dynamics can be described in two physically equivalent representations:

1. *The Unconstrained Representation:* The system is expressed in the full N-residue basis, retaining the centre-of-mass degree of freedom.
2. To prevent non-physical translational drift, the driving noise is restricted to the internal fluctuation subspace using the projection operator 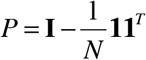, (where **I** is the *N* × *N* identity matrix and **1** is the all-ones vector) yielding a noise covariance proportional to *P*.
3. *The Center-Fixed Subspace Representation:* Global center-of-mass motion is projected out initially using *P*, restricting dynamics purely to internal shape deformations where *P* acts as the identity operator within the fluctuation subspace.

Solving the Langevin equation yields the time-delayed covariance matrix for a lag time *τ* :

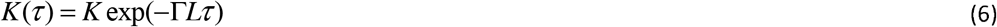

The derivation of the overdamped Langevin equation, together with the identity *LK* = *KL* = *P* on the fluctuation subspace, is given in Appendix A.

For small time delays *τ*, expanding the matrix exponential yields the short-time covariance elements. In the reduced subspace representation, the leading temporal corrections read:

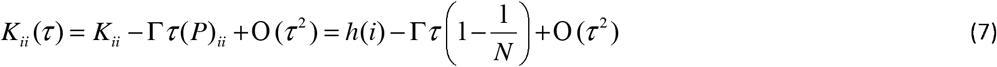

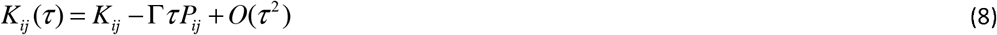

For contacting off-diagonal pairs (*i* ≠ *j*), *K*_*ij*_ (*τ*) acquires the leading finite-size translational term 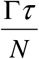 from the center-of-mass projector *P*, which must be retained in finite-*N* information-theoretic expansions.

### 2.3 Transfer Entropy and Exact Factorization of Asymmetry

Transfer entropy quantifies the directed information flow between stationary processes. For a continuous multivariate Gaussian process, the differential entropy of an *m* -dimensional variable with covariance matrix **Σ** is given by 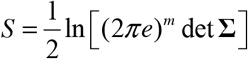. The transfer entropy *T*_*i*→ *j*_ (*τ*) measures the reduction in uncertainty of the future state *x*_*j*_ (*t* +*τ*) provided by the present state *x*_*i*_ (*t*), conditioned on the present state *x*_*j*_ (*t*) :

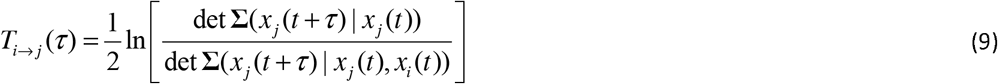

Using Schur complements of the partitioned joint covariance matrices, conditional variances reduce analytically to ratios of submatrix determinants [19, 20]. The short-time expansion of these determinant ratios, and the cancellation of the Laplacian degree terms in their difference, are carried out in Appendix B.

The net directional communication between residues *i* and *j* is evaluated via the transfer entropy asymmetry:

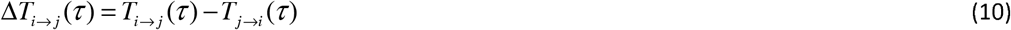

Expanding the determinant ratios for short time delays *τ* → 0^+^, the diagonal degree terms of the Laplacian matrix *L* cancel identically in the difference. This cancellation is exact and requires no structural approximation beyond the Taylor expansion.

To leading order in time delay *τ*, the net transfer entropy asymmetry factors into the exact product form:

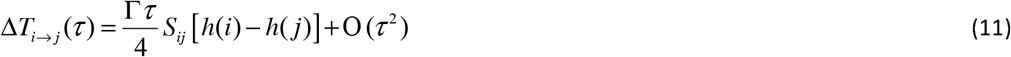

where *S*_*ij*_ is a symmetric structural coupling factor defined by:

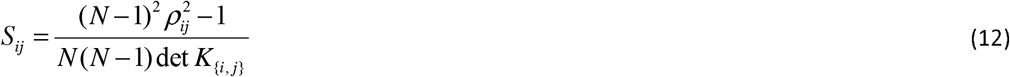

Its sign is determined by *S*_*ij*_ 0 ⟺| *ρ*_*ij*_ |> 1 / (*N* −1) and det 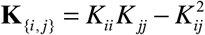 is the determinant of the pairwise covariance submatrix, and 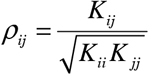 is the normalized Pearson correlation coefficient.

Equation (11) represents the central analytical result of this paper: the direction of net information transfer is determined strictly by the spatial gradient of the scalar node potential, ∇*h*(*i, j*) = *h*(*j*) − *h*(*i*), while the symmetric coupling factor *S*_*ij*_ modulates its magnitude. For all physical contact pairs satisfying 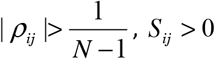, ensuring that information flows strictly from high-potential source residues (large *h*(*i*), high flexibility) toward low-potential sink residues (small *h*(*j*), high rigidity).

### 2.4 Normalized Field, Integrability, and Path Independence

For pairs with non-zero structural coupling (*S*_*ij*_ > 0), we define the normalized short-time asymmetry as:

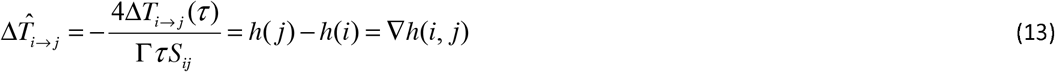

The normalized asymmetry field 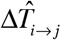 is strictly antisymmetric 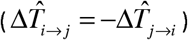 and represents a conservative discrete gradient field defined on the network nodes.

Consequently, for any path P = {*v*_1_, *v*_2_,⃛, *v*_*m*_ } connecting residue *v*_1_ to residue *v*_*m*_, the accumulated normalized asymmetry telescopes identically:

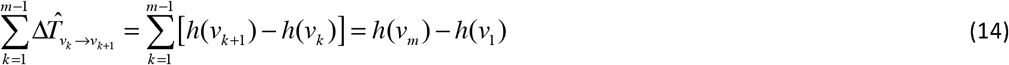

This proves that short-time normalized information transfer is strictly path-independent. Around any closed cycle C on the residue interaction network, the circulation vanishes:

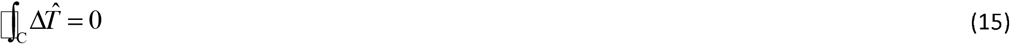

Information flow in the short-time regime is therefore purely irrotational and potential-driven. When two residues occupy an equipotential state, *h*_*i*_ = *h*_*j*_, the first-order directional driving force vanishes identically. As shown in Appendix B, the directional asymmetry then generically begins at *O*(*τ* ^2^) : its direction is determined by the local weighted-coordination difference *L*_*ii*_ − *L*_*jj*_, while its magnitude is modulated by the second-order pair geometry through *S*_*ij*_ (or equivalently, *D*_*ij*_ or *R*_*ij*_).

### 2.5 Sign Persistence: Extending Potential Gradients Beyond Short Times

While the expansion in Eq. (11) establishes the initial directional tendency as *τ* → 0^+^, physical information processing in proteins occurs over finite relaxation windows. We define the sign-persistence principle for equilibrium linear Gaussian dynamics: across the physical relaxation spectrum 0 < *τ* ≤ *τ*_relax_, the directional asymmetry Δ*T*_*i*→ *j*_ (*τ*) maintains a constant sign matching its short-time gradient:

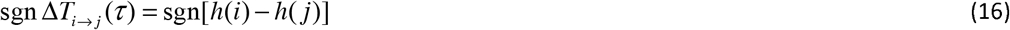

Sign persistence is an empirical regularity of equilibrium relaxation rather than a general theorem; its formal statement, and the propositions establishing that the integrated transfer and the peak comparison inherit the short-time direction, are given in Appendix D. As we show below, 96.86% of the 796 KRAS contacts retain their short-time sign over 0.05 ≤ τ ≤ 5.0, and 3.14% reverse at long lag (Supplementary Section S7).

Sign persistence ensures that the short-time analytical potential gradient ∇*h*(*i, j*) generalizes directly to finite-time and time-integrated communication metrics:

Cumulative Integrated Transfer: The total net information transfer over a finite relaxation window [0,*τ*_max_], defined as 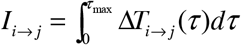, accumulates without destructive cancellation. Its sign is guaranteed to agree with the node potential difference: sgn *I*_*i*→ *j*_ = sgn[*h*(*i*) − *h*(*j*)]. Peak Transfer Comparison: For a sign-persistent pair with h(i)>h(j), and then

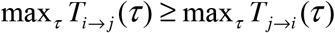

Sign persistence converts the short-time gradient ∇*h*(*i, j*) into a robust, global predictor of finite-time directional information flow across the protein fold.

### 2.6 The Experimental B-Factor Direction Map

By fluctuation-dissipation theorem, the physical covariance matrix is related to the topological pseudoinverse *K* by 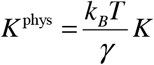, where *γ* is the effective network spring constant. The theoretical isotropic B-factor predicted by the Gaussian Network Model for residue *i* is:

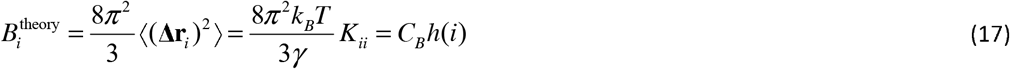

where *C*_*B*_ is a positive global scale factor.

Because experimental atomic displacement parameters reported in PDB files 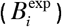 reflect local spatial variance, 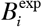 serves as an empirical direct proxy for *h*(*i*). We define the experimental B-factor direction matrix as:

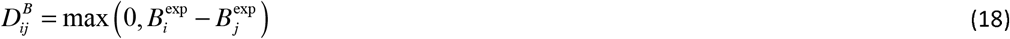

An oriented edge *i* → *j* exists whenever 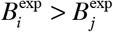.

This experimental B-factor mapping provides a fitting-free, parameter-free directional baseline: the directional source-and-sink topology of any protein fold can be evaluated directly from experimental *B* - factors without assuming kinetic rate constants, specifying time lags, or selecting an explicit network cutoff distance *r*_*c*_.

## 3. Results and Structural Analysis

### 3.1 Parameter-Free Communication Landscape of Oncogenic KRAS

To demonstrate the dynamic potential framework, we apply our analytical formulation to active-state human KRAS (PDB ID: 6GOD, chain A), establishing the baseline dynamic landscape that governs oncogenic KRAS signaling. The residue interaction network is constructed using *C*_*α*_ coordinates across the 172 resolved residues. A weighted Kirchhoff matrix *L* is constructed using C_*α*_ coordinates across the 172 resolved residues with a contact cutoff of *r*_*c*_ = 7.8 *Å* and a distance-dependent exponential weighting *γ*_*ij*_ = exp(−*d*_*ij*_ / *T*) with *T* = 1 (following our previous work [16]), yielding a connected graph of 796 weighted edges. The complete set of fixed model parameters and the principal numerical checks are listed in Supplementary Table S1.

The Moore–Penrose pseudoinverse *K* = *L*^+^ is evaluated directly from the weighted contact topology. The diagonal elements *h*(*i*) = *K*_*ii*_ define the local residue fluctuation potential, representing the variance in reduced units. The pairwise transfer entropy asymmetry in the short-time regime takes the exact factored form 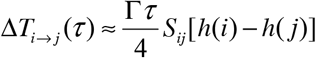, where *S*_*ij*_ is the non-negative structural coupling factor.

Figure 1 displays the integrated directional transfer entropy asymmetry map, 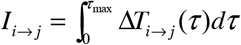, capturing the cumulative net information flow between residue pairs across the relaxation window. Because short-time directional gradients persist across extended delay times via the sign-persistence principle, the integrated transfer map retains the underlying potential-driven topology while accounting for finite-time relaxation.

**Figure 1.**
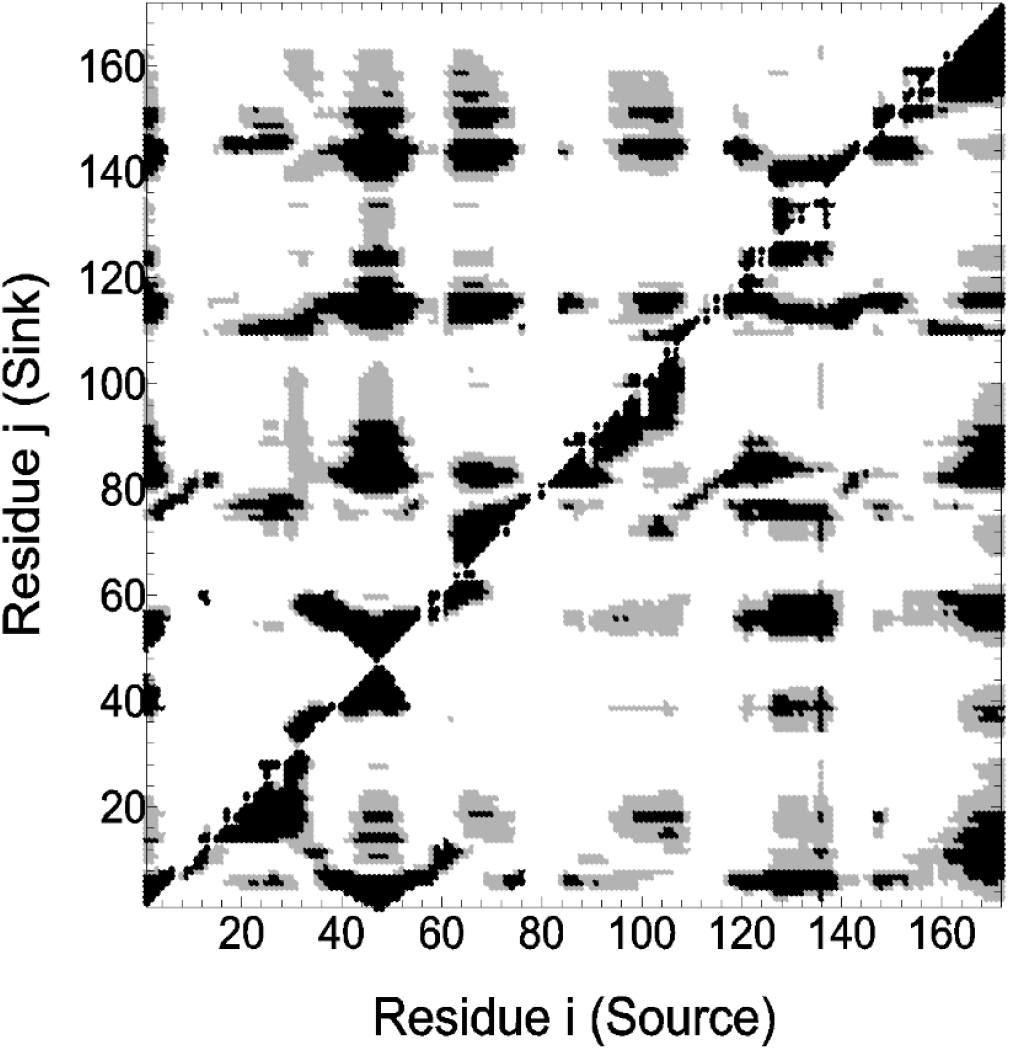
Integrated directional entropy-transfer asymmetry map for KRAS (PDB: 6GOD). The horizontal axis represents the source residue *i*, and the vertical axis represents the sink residue *j*. Interaction pairs are grouped into four equal-frequency quartiles. Gray points highlight pairs in the 3rd quartile (50th–75th percentile range), while black points indicate pairs in the top quartile (75th–100th percentile of directional transfer magnitude). Net directionality closely tracks the fluctuation potential gradient Δ*h*(*i, j*) = *h*(*i*) − *h*(*j*). The elastic network is constructed from the C_*α*_ coordinates of chain A (172 residues) using a cutoff distance of *r*_*c*_ = 7.8 *Å* and exponential distance weighting (*T* = 1), yielding 796 weighted contacts.

For every contacting pair in KRAS, the normalized pairwise correlation satisfies 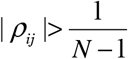, fixing the net directional information flow from flexible regions (large *h*(*i*)) toward rigid regions (small *h*(*j*)). Finite-delay calculations distinguish two related but different measures of directional consistency. Among the 796 weighted contacts, 96.86% retain their short-time sign throughout the sampled interval 0.05 ≤ *τ* ≤ 5.0. Over all 14, 706 residue pairs, the short-time sign agrees with the sign of the integrated asymmetry *I*_*i*→ *j*_ in 98.81% of cases (Supplementary Section S7).

Summing the integrated transfer over immediate network contacts yields the net residue-level flux balance, 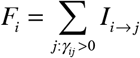. The sum runs over the immediate network contacts of residue i. Net information sources (*F*_*i*_ > 0) concentrate in flexible loops and terminal segments (e.g., Thr148, Gln25, Gly48, and Asp47), whereas major information sinks (*F*_*i*_ < 0) reside within tightly coordinated structural elements, including Switch II (Gly60) and core helices (Ser145, Gln22, Leu52, and Ala134). The single strongest integrated pairwise contact asymmetry in the entire network occurs at the Gln61 → Gly60 junction (*I*_61→60_ = 1.841 *nats*), confirming that Gln61 acts as a primary dynamic source transmitting persistent memory directly into the catalytic sink at Gly60. Notably, this theoretical source-sink relationship directly mirrors experimental crystallographic displacement parameters in PDB 6GOD, where the *C*_*α*_ *B* -factor drops steeply from 47.73*Å*^2^ at Gln61 down to 27.68*Å*^2^ at Gly60.

### 3.2 Reading Transfer Maps Through the Laplacian Minor Hierarchy

The complex communication channels highlighted in the transfer entropy heatmaps are structured through the algebraic minors of the covariance matrix *K*. Rather than treating the minor hierarchy as a separate geometric concept, we employ it here as the natural multi-body structural language required to explain the bandwidth, pathway overlap, and hub organization of entropy transfer:

1. *Order-1 Minor (Node Potential h*(*i*) - *Fluctuation Driving Force):* The single-node diagonal element determines the local variance and conformational entropy. Spatial potential differences ∇*h*(*i, j*) = *h*(*j*) − *h*(*i*) provide the fundamental driving force for directional asymmetry.
2. *Order-2 Minors: Effective Distance and Pairwise Channel Bandwidth*. While the scalar node potential *h*(*i*) = *K*_*ii*_ represents the foundational order-1 minor (local variance), the order-2 minors of the covariance matrix govern pairwise communication. Specifically, the effective resistance distance *R*_*ij*_ = *h*(*i*) + *h*(*j*) − 2*K*_*ij*_ measures the inverse capacity, or spatial channel bandwidth, of signal propagation between nodes *i* and *j*. A low effective distance *R*_*ij*_ indicates a high-bandwidth coupling mediated by multiple redundant physical pathways, whereas a high *R*_*ij*_ reflects a restricted, low-bandwidth communication bottleneck.
3. *Order-3 Minors: Triad Cooperation and Integrated Allosteric Sectors:* While order-2 minors (*R*_*ij*_) characterize pairwise communication channels, order-3 minors, determinants of 3×3 covariance submatrices det(*K*_{*i, j,k*}_), quantify multi-body joint fluctuations across three-residue triads. To assess whether signal propagation from a central source branches into independent channels or operates through a shared structural manifold, we define an order-3 triad cooperation index via the normalized overlap of joint determinants:

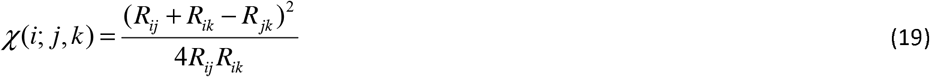 This index is the squared cosine between the effective displacement vectors from residue i toward j and k, and is bounded by 0 ≤ χ ≤ 1 by the Cauchy–Schwarz inequality; the derivation is given in Appendix C. Applied to KRAS, this metric yields a high normalized overlap for the {Gln61, Gly60, Phe156} triad, indicating that signal transmission from Switch II (Gln61) to the catalytic pocket (Gly60) and the distal core (Phe156) converges on a single, mechanically integrated communication sector rather than routing through decoupled, independent pathways. This network interpretation is strongly supported by functional and dynamical studies of KRAS. Switch-II motions have been shown to drive long-range dynamics across both Switch I and distal allosteric elements (Vatansever et al., 2016 [21]; Volmar et al., 2022 [22]). Furthermore, deep mutational scanning demonstrates that substitutions at Gly60 and Phe156 broadly perturb effector-binding landscapes across multiple interaction partners (Weng et al., 2024 [23]), confirming that these specific residues belong to a common, functionally sensitive allosteric network (Sethi et al., 2009 [24]).
4. Order-4 Minors: Submatrix Determinants and Multi-Node Relay Hubs. Beyond single-node potentials (1×1) and pairwise channels (2×2), higher-order minors capture multi-body collective correlations. Specifically, an order-4 minor, the determinant of a 4×4 submatrix of the covariance matrix *K*, measures the four-body joint fluctuation volume det(*K*_{*i, j,k,l*}_). Geometrically, this determinant quantifies the spatial volume of collective uncertainty shared by four residues; a compact joint volume indicates that the group’s thermal motions are tightly constrained to a low-dimensional manifold. In structural networks, order-4 minors identify critical relay nodes: residues situated at steep local potential gradients Δ*h* that dynamically link distinct functional domains. In our KRAS network, Phe156 provides a clear example of such a relay node. Although it acts as a rigid structural sink with a small local potential *h*(Phe156), its 4-body joint fluctuations remain tightly correlated with both the active-site switch loops and distal helical elements. This places Phe156 in a strategic communication pathway linking the effector lobe (residues 1–86) and allosteric lobe (residues 87–166). This observation is strongly consistent with prior work demonstrating that long-range directional coupling between the active site and allosteric core is mediated through the *α* 3 –L7 region and helices *α* 4 –*α* 5 (Vatansever et al., 2016 [21]; Volmar et al., 2022 [22]), as well as deep mutational studies establishing Phe156 (*α* 5) as a distal allosteric hotspot influencing effector binding (Weng et al., 2024 [23]).

### 3.3 Sequence Distance Regimes and Experimental Validation

Filtering the network asymmetry by sequence separation shows two distinct functional regimes:

- *Local Chain Dynamics (* | *i* − *j* |≤ 3): Directional transfer is dominated by nearest-neighbor sequence contacts along the covalent backbone, accounting for 78 of the 100 strongest pairwise asymmetries due to strong local spatial covariance.
- *Nonlocal Allosteric Channels (* | *i* − *j* |> 3): Nonlocal interactions show long-range communication corridors across the tertiary fold. Nonlocal directional flux concentrates into a specific spatial corridor connecting Switch I (residues 39–46) directly to Switch II (residues 51–56), anchored by key inter-residue directional pairs such as Val45 → Cys51 and Ile46 → Cys51.

To validate the theoretical model against independent physical observables, the analytical potential profile *h*(*i*) was compared with experimental crystallographic B-factors from PDB 6GOD. The theoretical potential correlates strongly with experimental B-factors (Spearman *r*_*s*_ = 0.807). Furthermore, the direction of net information transfer calculated analytically from Eq. (11) agrees with the experimental B-factor gradient 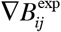 for 77.6% of all weighted network contacts. These results indicate that asymmetric entropy transfer across structural gradients can be derived directly from experimental atomic displacement parameters without introducing empirical parameters or running molecular dynamics simulations. Controls for contact degree, conductance heterogeneity, and a degree-preserving rewired null model are reported in Supplementary Section S6.

This analytical framework directly unifies and formalizes the distinction between local residue mobility and directional information flow in thermal equilibrium [4, 8]. While local conformational variance, derived directly from experimental atomic displacement parameters via the order-1 node potential *h*(*i*), measures isolated spatial mobility, directional asymmetry is governed by conditional variance reduction across the potential gradient Δ*h*_*ij*_. Consequently, large local flexibility does not automatically imply high outgoing information transfer; a highly mobile surface loop may absorb thermal energy while remaining conditionally decoupled from downstream functional sites if structural coupling *S*_*ij*_ is weak. Conversely, relatively rigid core residues, characterized by small local potentials *h*(*j*), frequently act as structural entropy sinks or higher-order relay hubs (such as Phe156 in KRAS) that constrain the multi-body joint fluctuation manifold. By formalizing these conditional dependencies through the Laplacian minor hierarchy and Schur complements, our formulation provides a rigorous mathematical bridge between static experimental B-factor maps and collective allosteric communication pathways without requiring empirical parameter fitting or trajectory simulations.

Finally, comparing the transfer entropy flux with combinatorial Hodge decomposition shows that information flow in KRAS is overwhelmingly hierarchical [13]. The irrotational gradient component accounts for 98.2% of the total transfer entropy flux, while local 3-clique recirculation (curl) and cavity circulation (harmonic) carry less than 2% of the signal. The scarcity of the harmonic component confirms that information transfer does not loop or recirculate locally, but flows unidirectionally from high-potential source regions down a strictly irrotational gradient toward low-potential sink regions.

## 4. Discussion and Conclusion

The principal theoretical achievement of this work is the exact factorization of the short-time transfer entropy asymmetry Given by Eq. 11. This result establishes that directed information flow in an equilibrium linear Gaussian network is generated to leading order by the spatial discrete gradient of a scalar node potential, *h*. The symmetric, non-negative coupling factor *S*_*ij*_ modulates the numerical magnitude of the flux but leaves its sign and direction strictly governed by *h*_*i*_ − *h*_*j*_. This directionality aligns directly with recent multi-enzyme analyses by Miño-Galaz et al., [25] who demonstrated that entropy and information are systematically transferred from peripheral, highly flexible surface regions toward rigid, buried active sites across diverse enzymatic architectures.

An equipotential pair provides a useful limiting case. When *h*_*i*_ = *h*_*j*_, the two residues share the same equilibrium fluctuation amplitude, causing the first-order directional bias to disappear identically. Equal fluctuation amplitudes, however, do not imply identical local dynamics. The diagonal elements *L*_*ii*_ and *L*_*jj*_ measure the total weighted coupling of each residue to its structural environment, characterizing their respective local restoring constraints. If *L*_*ii*_ ≠ *L*_*jj*_, these constraints are unequal, and a directional asymmetry reappears at *O*(*τ* ^2^). The pair geometry, captured by *S*_*ij*_ or, equivalently, the effective resistance *R*_*ij*_, modulates how strongly this kinetic inequality is expressed.

Directionality therefore follows a natural hierarchy: differences in equilibrium fluctuation amplitude drive asymmetry at first order, whereas differences in local coordination govern transport when the fluctuation potential is degenerate. Crucially, this remains an equilibrium phenomenon; it reflects unequal conditional relaxation rather than a non-equilibrium thermodynamic current. Finally, if both *h*_*i*_ = *h*_*j*_ and *L*_*ii*_ = *L*_*jj*_, the quadratic contribution also vanishes, deferring directional discrimination to higher orders in time.

Several physical and structural implications of this framework warrant emphasis:

1. *Exactness of Degree Cancellation*: The degree terms cancel identically in the derivation of the short-time asymmetry. This cancellation is exact and holds for any connected graph with arbitrary positive edge weights, regardless of graph degree distribution or topological heterogeneity.
2. *Thermodynamic Interpretation of Directionality*: The directional asymmetry does not imply a non-equilibrium heat flux or thermodynamic dissipation. In accordance with equilibrium thermodynamic interpretation, the system remains in full thermodynamic equilibrium throughout. The observed asymmetry represents an intrinsic inequality in conditional predictability: knowledge of the present state of a flexible source residue (large local potential, persistent fluctuation memory) reduces uncertainty about the future state of a rigid sink residue (small local potential, fast relaxation) to a greater degree than the reverse.
3. *Physical Basis of Sign Persistence:* The sign-persistence principle asserts that the initial direction dictated by the short-time gradient remains invariant over physical relaxation windows. While direction reversals can theoretically occur in chaotic or delay-coupled non-linear systems, linear response relaxation in overdamped macromolecular networks suppresses high-frequency mode crossovers, ensuring that short-time gradients reliably predict finite-time cumulative integrals and peak transfer magnitudes.
4. *The Experimental B-Factor Laboratory:* Because the theoretical node potential *h*(*i*) is directly proportional to mean-square atomic displacement, classical equipartition bridges theoretical covariance directly to experimental crystallographic *B* -factors: 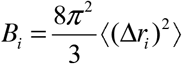. This equivalence converts high-quality structural deposits in the Protein Data Bank into an immediate, parameter-light empirical laboratory. Without empirical parameter fitting, force-constant tuning, or MD trajectory sampling, the directional source-and-sink topology of any protein fold can be read directly from standard ATOM records in a PDB file.
5. *Role of the Laplacian Minor Hierarchy*: By placing *h*(*i*) as the Order-1 floor of the Laplacian minor hierarchy, we establish a systematic foundation for multi-body allosteric theory. When two residues occupy an equipotential state, *h*(*i*) = *h*(*j*), the *O*(*τ*) pairwise directional term vanishes. This does not, however, imply a progression to successively higher covariance minors. Direct expansion shows that the leading pairwise asymmetry then generally occurs at *O*(*τ* ^2^), with its direction determined by the local weighted-coordination difference *L*_*ii*_ − *L*_*jj*_ and its magnitude modulated by the pair geometry through *S*_*ij*_, equivalently *R*_*ij*_ or *D*_*ij*_. Order-three and higher invariants instead characterize genuinely many-body structural relations and enter naturally when additional residues are included explicitly in conditioned or collective observables. Thus, the covariance-minor hierarchy and the short-time expansion constitute distinct organizations of the theory: the former is a hierarchy in the number of residues involved, whereas the latter is a hierarchy in powers of the delay time*τ*.
6. *Future Perspectives: Partial Correlations and Conditional Independence*: The linear-algebraic framework established here lays the foundation for systematic conditional independence testing in macromolecular fluctuation networks. In Gaussian graphical models, zero entries in the concentration matrix (*L*_*ij*_ = 0) denote conditional independence given the remainder of the system, allowing direct mechanical couplings to be distinguished from indirect, multi-step correlations. Extending our Laplacian minor hierarchy to evaluate high-order partial correlations (*ρ*_*ij* S_) via Schur complements will enable systematic network pruning, isolating direct communication backbones from background thermal noise. Furthermore, quantifying conditional decoupling across submatrices can identify obligatory allosteric “choke points”, intermediate relay nodes whose conditional shielding completely decouples distant functional domains. Applied to multi-state structural ensembles and cryo-EM landscapes, this extension promises a parameter-free approach for predicting driver mutation effects and discovering latent drug-targetable sites directly from experimental atomic coordinates.

In conclusion, by unifying linear response theory, elastic network statistics, and transfer entropy within the algebraic structure of the Laplacian minor hierarchy, this work establishes a rigorous, parameter-free foundation for macromolecular directional communication.

## Data availability

The Supplementary archive contains the parameter summary, the 796 contact values, the residue table listing *h* (*i*), the standardized B-factor, *F*_*i*_, the uniform-conductance control and contact degree, the 1000 null replicates, the supplementary figure files, the analysis script, and the full 14,706-pair report. An audio podcast providing an overview of the core concepts and results of this work is available on Zenodo at https://doi.org/10.5281/zenodo.21909209

## Appendices

These four appendices collect the derivations used in the main text. The notation is fixed throughout. *L* is the weighted Kirchhoff matrix built from the positive pair weights *γ*_*ij*_ ; *K* = *L*^+^ is its Moore–Penrose pseudoinverse; Γ is the relaxation-rate prefactor; and *N* is the number of residues. The projector orthogonal to the translational zero mode is

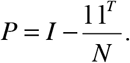

The residue fluctuation potential is *h* (*i*) = *K*_*ii*_, the squared effective distance is

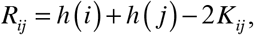

and, for a fixed pair,

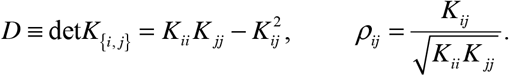

The symmetric structural coupling factor *S*_*ij*_ is defined in Eq. (B16). Directional quantities use the notation *T*_*i*→*j*_ (*τ*), Δ*T*_*i*→*j*_ (*τ*) = *T*_*i*→*j*_ (*τ*) − *T*_*j*→*i*_ (*τ*), ∇*h* (*i, j*) = *h* (*j*) − *h* (*i*), *I*_*i*→ *j*_ = ∫ Δ*T*_*i*→ *j*_ (*τ*) *dτ*, and 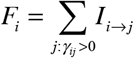. The sum runs over the network contacts of residue *i*. We write *u* = Γ*τ* for the reduced delay. The transfer entropy is evaluated for a single Cartesian displacement coordinate. Since the Gaussian network is isotropic, the three Cartesian components are equivalent. Natural logarithms are used, and the entropy is therefore measured in nats.

The pair weights are always denoted by *γ*_*ij*_. The scalar *γ* used in Section 2.6 is the effective network spring constant and is kept separate from the pair weights. The scalar *T* in the distance-weighting function never denotes transfer entropy, which always carries a directional subscript.

## Appendix A. Langevin Derivation of the Time-Dependent Covariance

This appendix derives the delayed covariance *K* (*τ*) = *e*^−Γ*Lτ*^ *K* and the identity *LK* = *KL* = *P*.

### A.1 The overdamped Langevin equation

For one Cartesian displacement component, the overdamped network dynamics are

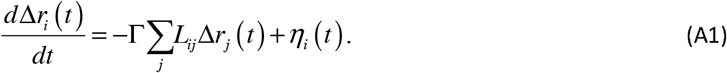

In matrix form, this is

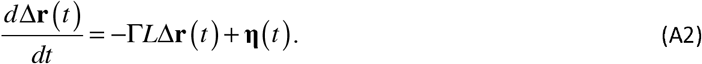

Here Γ has dimensions of an inverse time. It is a relaxation-rate prefactor, not a friction coefficient. In reduced fluctuation units, the noise has zero mean and covariance

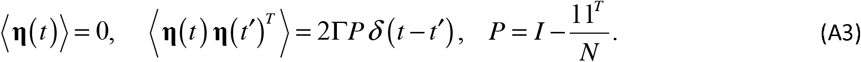

The projector *P* removes the translational zero mode and prevents unbounded center-of-mass drift. The stationary covariance Σ satisfies the Lyapunov equation

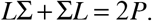

This equation alone does not fix the zero mode: *K* + *α*11^*T*^ is also a solution. The center-fixed ensemble imposes

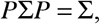

which is equivalent to suppressing fluctuations of the center-of-mass mode. This condition removes the zero-mode freedom. Since *LK* = *KL* = *P*, the unique centered stationary covariance is Σ= *K* = *L*^+^. Restoring the physical spring constant *γ* and thermal energy *k*_*B*_*T* gives

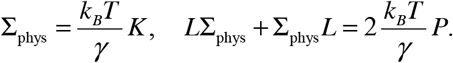

Equivalently, the physical noise covariance is 2Γ(*k*_*B*_*T* / *γ*) *Pδ* (*t* − *t*′). This is the fluctuation–dissipation relation in the present normalization.

### A.2 Solution and the time-delayed covariance

The linear equation (A2) is solved by variation of constants:

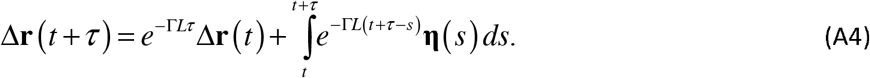

For *τ* > 0, the stochastic integral contains only noise after time *t* and is therefore uncorrelated with Δ*r* (*t*). Multiplying Eq. (A4) by Δ*r* (*t*)^*T*^, averaging, and using stationarity gives

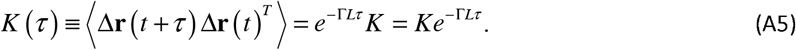

The last equality holds because *L* and *K* = *L*^+^ share the same orthonormal eigenvectors and therefore commute.

### A.3 The identity *LK* = *KL* = *P*

For a connected network, *L* is symmetric and positive semidefinite with null space spanned by the constant vector 1. The Moore–Penrose inverse acts as the ordinary inverse on the orthogonal complement of that null space and vanishes on the null space. Hence

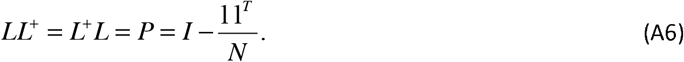

Thus *LK* = *KL* = *P*. On the subspace orthogonal to 1, *P* acts as the identity. In particular, *K*1 = 0 and *PKP* = *K*.

### A.4 Short-time expansion

Set *u* = Γ*τ*. A Taylor expansion of the matrix exponential gives

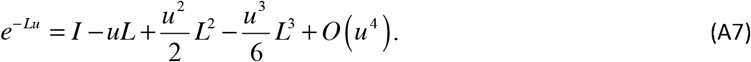

Multiplying Eq. (A7) on the right by *K* converts each power of *L* into a lower power through *LK* = *P, L*^2^*K* = *LP* = *L*, and *L*^3^*K* = *L*^2^. Therefore

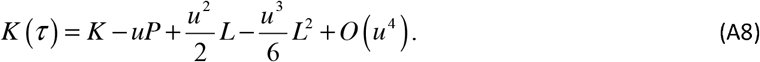

Equations (A9) and (A10) are now obtained simply by taking matrix elements of Eq. (A8). For a diagonal element, *P*_*jj*_ = 1−1/ *N* ; keeping terms through *u*^*2*^ gives

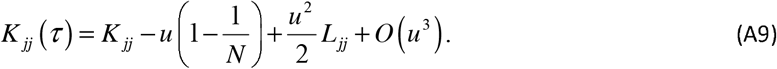

For an off-diagonal element, *i* ≠ *j* and *P*_*ij*_ = −1 / *N*. The term −*uP*_*ij*_ therefore becomes +*u* / *N*, so

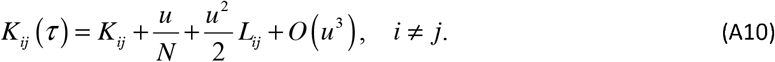

Thus the diagonal correction begins at order *u*. The off-diagonal correction also contains an order-u term, *u* / *N*, produced entirely by the center-of-mass projector. This finite-N term must be retained in the transfer-entropy expansion.

## Appendix B. Short-Time Expansion of the Transfer Entropy Asymmetry

This appendix derives Eq. (11) of the main text. The calculation shows that the directional transfer entropy is linear in delay, that the Laplacian degree terms cancel from the leading asymmetry, and that the remaining direction is set by the fluctuation-potential difference.

### B.1 Deviations of the covariance elements

Let *j*^+^ = *x*_*j*_ (*t* +*τ*), while *i* = *x*_*i*_ (*t*) and *j* = *x*_*j*_ (*t*). Equations (A9)–(A10) give the delayed covariance as its equal-time value plus a small correction. It is convenient to isolate those corrections:

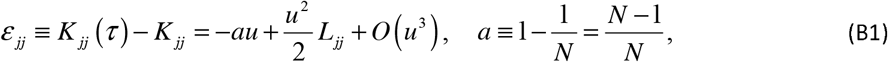

and

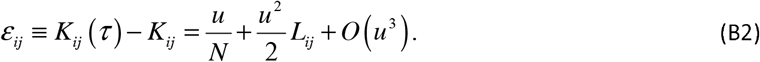

The covariance matrices needed for the two conditional variances are therefore

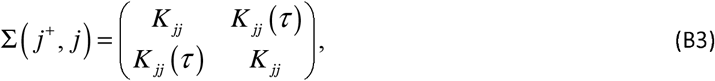

and

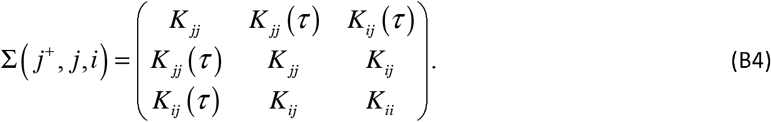

We consider nondegenerate pairs with

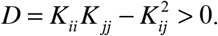

### B.2 The two conditional variances

For jointly Gaussian variables, conditioning is a Schur complement. Applied first to the 2 ×2 matrix in Eq. (B3), this gives

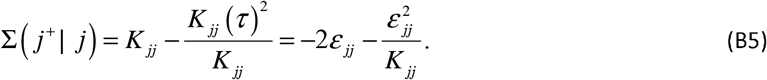

Substituting Eq. (B1) into Eq. (B5) and retaining terms through *u* ^2^ gives

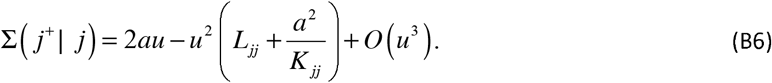

For conditioning on both present variables (*j, i*), let *c* = (*K*_*jj*_ (*τ*), *K*_*jj*_ (*τ*))^*T*^ and let *M* be their 2 ×2 equal-time covariance block. The Schur complement is

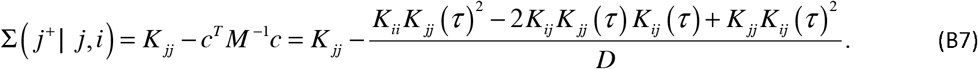

To pass from Eq. (B7) to Eq. (B8), substitute *K*_*jj*_ (*τ*) = *K*_*jj*_ + *ε*_*jj*_ and *K*_*ij*_ (*τ*) = *K*_*ij*_ + *ε*_*ij*_ and expand the numerator. The constant term is *K*_*jj*_ *D* and cancels the leading *K*_*jj*_ outside the fraction, as required because *j*^+^ = *j* at zero delay. The terms linear in *ε*_*ij*_ cancel as well. What remains is

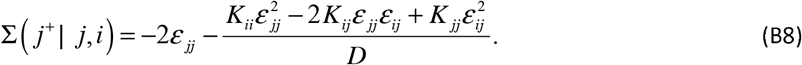

Now insert Eqs. (B1)–(B2). Since each *ε* is *O* (*u*), only their leading *O* (*u*) parts are needed inside the quadratic numerator of Eq. (B8); their *O* (*u*^2^) parts would contribute only at *O* (*u*^3^). This yields

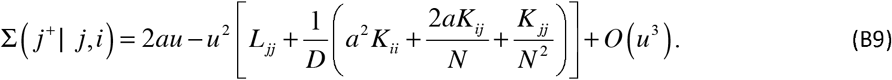

Both conditional variances begin with the same term, 2*au*. Their ratio therefore differs from unity first at order *u*.

### B.3 The log-ratio at order *τ*

*F*or Gaussian variables, transfer entropy is one half of the logarithm of the ratio of the two conditional variances:

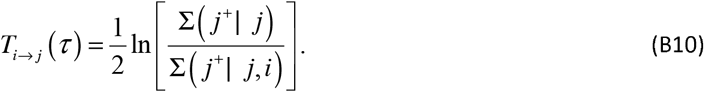

Factor the common leading term 2*au* from Eqs. (B6) and (B9). The ratio is then of the form (1+ *xu*) / (1+ *yu*), and ln [(1+ *xu*) / (1+ *yu*)] = (*x* − *y*)*u* + *O* (*u*^2^). This gives

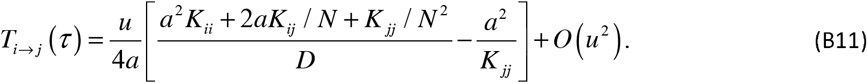

The term *L*_*jj*_ has disappeared because it entered both conditional variances with the same coefficient. To simplify the remaining covariance terms, put the bracket in Eq. (B11) over the common denominator *K*_*jj*_ *D* and use *a* = (*N* −1) / *N*. The factor 1/ *N* ^2^ must then be retained explicitly. The numerator is

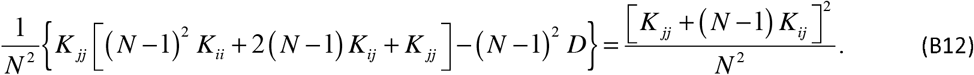

The factor 1/ *N* ^2^ in Eq. (B12), together with 1/ *a* = *N* / (*N* −1), gives the denominator *N* (*N* −1) below. With *u* = Γ*τ*,

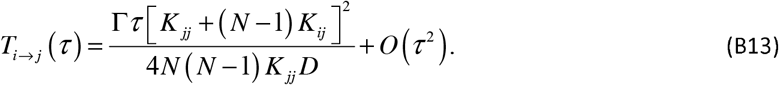

The leading term is nonnegative, as required. Interchanging *i* and *j* gives *T*_*j*→*i*_ (*τ*).

1 If *K*_*ij*_ = 0, Eq. (B13) gives *T*_*i*→ *j*_ (*τ*) = Γ*τ* / [4*N* (*N* −1) *K*_*ii*_] + *O* (*τ* ^2^). This small term is induced by the center-of-mass projector: *P*_*ij*_ = −1 / *N* produces the *u* / *N* contribution in Eq. (A10). It is a finite-size effect, of order *N* ^−2^ at fixed *K*_*ii*_, and does not represent direct pair coupling.

### B.4 The asymmetry at order *τ*

*T*he asymmetry is obtained by subtracting the expression in Eq. (B13) with *i* and *j* interchanged. The common positive prefactor may be taken outside the difference. The remaining algebra is

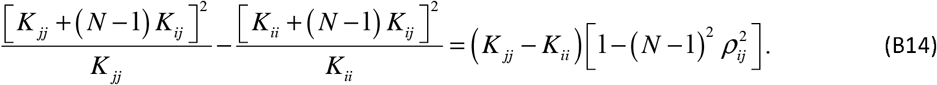

Since *K*_*jj*_ − *K*_*ii*_ = *h* (*j*) − *h* (*i*) = ∇*h* (*i, j*), Eq. (B14) separates the directional part from a symmetric pair factor. Hence

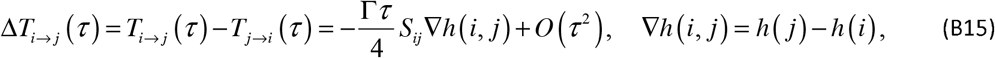

with

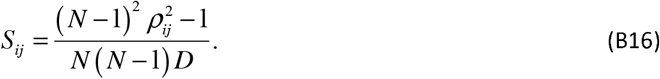

Equations (B15)–(B16) are Eqs. (11)-(12) of the main text. Their immediate consequences are: Exact cancellation. The diagonal degree terms *L*_*ii*_, *L*_*jj*_ and the off-diagonal term *L*_*ij*_ do not appear in the leading asymmetry. This cancellation holds for any connected graph with positive edge weights. Sign. *S*_*ij*_ is symmetric and *S*_*ij*_ > 0 exactly when |*ρ*_*ij* | *>*_ 1/ (*N* −1). For *N* = 172, the threshold is 5.85×10^−3^. The smallest |*ρ*_*ij*_| among the 796 KRAS contacts is 1.05×10^−2^ ; thus every contact lies above the threshold. For these pairs, *h* (*i*) > *h* (*j*) implies Δ*T*_*i*→*j*_ > 0 at sufficiently small positive delay. Threshold. *S*_*ij*_ = 0 when |*ρ*_*ij*_| = 1/ (*N* −1). A nonzero covariance *K*_*ij*_ alone therefore does not guarantee a nonzero leading asymmetry. If *S*_*ij*_ < 0, the leading direction is reversed. No general statement about the magnitude follows from the sign of *S*_*ij*_ alone.

If *h* (*i*) = *h* (*j*) with *S*_*ij*_ > 0, the linear term vanishes and the next term is derived in B.7. If *S*_*ij*_ = 0, the linear term also vanishes, but the next nonzero term must be determined separately; B.7 does not treat that case.

### B.5 Large-N consistency

Define

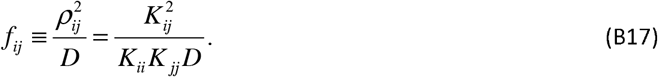

Substituting this definition into Eq. (B16) and separating the term generated by the projector gives

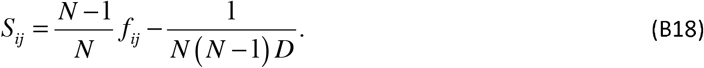

Thus *S*_*ij*_ tends formally to *f*_*ij*_ as *N* becomes large at fixed *K*. This statement only isolates the finite-N projector correction; it is not an asymptotic claim for a growing protein, because *K* itself changes with the network.

### B.6 Range of validity

Equation (B15) describes the initial Gaussian relaxation. The expansion requires Γ*τλ*_*m*_ □ 1 mode by mode, and Γ*τλ*_max_ □ 1 uniformly. In the Supplementary calculations Γ = 1. For KRAS at *τ* = 0.01, *τλ*_max_ = 1.29 × 10^−3^, well inside this regime. The expansion was also checked directly against the exact result. At *τ* = 10^−4^, 10^−3^, and 10^−2^, comparison of Eq. (B10) with Eq. (B15) gives a relative deviation proportional to *τ*, as required by the omitted *O* (*τ* ^2^) term; Eq. (B13) behaves likewise. Finite-delay behavior is treated in Appendix D.

### B.7 Equipotential pairs and the quadratic asymmetry

If *h* (*i*) = *h* (*j*), the coefficient in Eq. (B15) vanishes exactly:

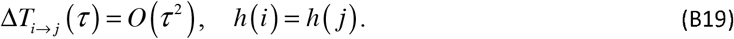

Write *h* (*i*) = *h* (*j*) = *h, c* = *K*_*ij*_, *D* = *h*^2^ − *c*^2^, and assume the pair is nondegenerate with *S*_*ij*_ > 0. Because each conditional variance starts at order *u*, the coefficient of *u* ^2^ in the transfer entropy depends on the conditional variances through order *u*^3^. We therefore retain one more term from Eq. (A8):

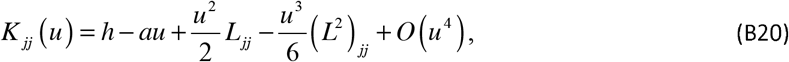

and, using *P*_*ij*_ = −1 / *N* for *i* ≠ *j*,

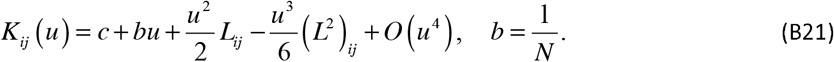

The expression for *K*_*ii*_ (*u*) follows from Eq. (B20) by interchanging *i* and *j*. Write

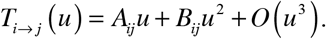

Substituting Eqs. (B20)–(B21) into the same Schur-complement formula used above and expanding the logarithm gives a simpler cancellation pattern. In *T*_*i*→ *j*_, the 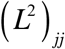 contribution cancels between its two conditional variances; the corresponding (*L*^2^)_*ii*_ term cancels within *T*_*j*→ *i*_. The term (*L*^2^)_*ij*_ first enters the conditional variance at order *u* ^4^ and therefore does not appear here. The remaining *L*_*ij*_ contribution is symmetric under *i* ↔ *j* and cancels only when the two directions are subtracted. The result is

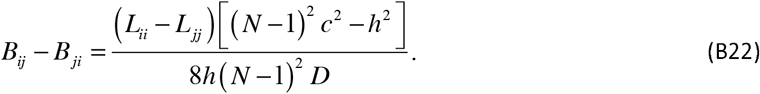

On the equipotential manifold, Eq. (B16) becomes

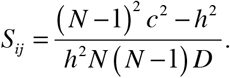

Using this identity to replace the covariance factor in Eq. (B22), and recalling *u* = Γ*τ*, gives

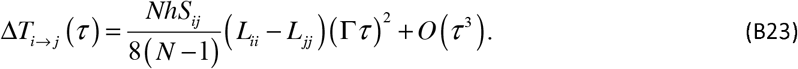

Since *S*_*ij*_ > 0 and *h* > 0, the prefactor in Eq. (B23) is positive. The sign is therefore selected only by the coordination difference:

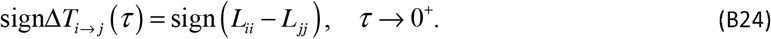

For a weighted Kirchhoff matrix,

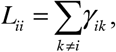

so the quadratic direction is fixed by the difference in local weighted coordination. If *L*_*ii*_ = *L*_*jj*_, the quadratic coefficient also vanishes and the first nonzero term occurs at *O* (*τ* ^3^) or beyond. Thus an equipotential pair is generically, but not unconditionally, quadratic.

The coefficient also has a simple geometric form. On the equipotential manifold, the definition of *R*_*ij*_ gives *R*_*ij*_ = 2(*h* − *c*). Solving this relation for *c* and substituting into *D* = *h*^2^ − *c*^2^ yields

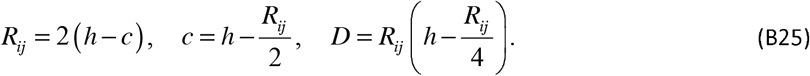

Substituting Eq. (B25) into Eq. (B16) eliminates *c* and *D* in favor of the pair geometry:

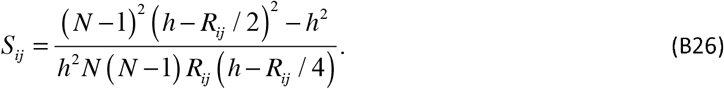

The magnitude of the quadratic response is set by the pair geometry, while its sign is selected by *L*_*ii*_ − *L*_*jj*_. The coefficient closes on *h, R*_*ij*_ (equivalently *D*), *S*_*ij*_, and the coordination difference. No order-three invariant enters at this order.

The linear and quadratic branches compare different quantities. Away from equipotentiality, the leading term compares equilibrium fluctuation amplitudes. On the equipotential manifold, the surviving term compares local relaxation constraints. Equal variance does not imply equal short-time dynamical response.

## Appendix C. The Fluctuation Hilbert Space and the Effective Distance

This appendix gives the geometric interpretation of *K*, shows that *R*_*ij*_ is a squared distance, derives the corresponding inner product, and defines the three-residue cooperation index used in the main text.

### C.1 The internal fluctuation space

The fluctuation geometry is the geometry induced by *K* : it describes how residues are separated, aligned, and coupled by their thermal fluctuations rather than by their Cartesian positions [3]. The weighted construction of *K* used here follows our earlier network formulation [16]. Geometry alone does not assign a direction; direction enters through the scalar potential difference *h* (*i*) − *h* (*j*).

The internal fluctuation space is

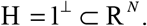

Each residue *i* is represented by *v*_*i*_ = *K*^1/2^*e*_*i*_ ∈H, with

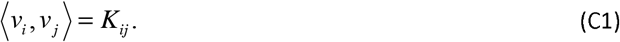

Thus *K* is the Gram matrix of the residue vectors. Setting *i* = *j* gives

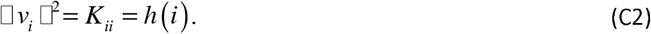

### C.2 The effective distance is a squared distance

Choose a reference residue *r* and define the relative Gram displacement

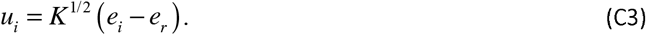

Using the definition of *v*_*i*_ in Eq. (C1), its inner product with *u*_*j*_ is

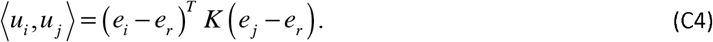

Setting *j* = *i* turns this inner product into a squared norm. Expanding the quadratic form then gives

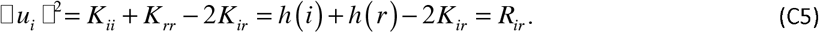

Hence *R*_*ij*_ is the squared distance between the Gram images of residues *i* and *j*. For a connected graph, *R*_*ij*_ ≥0, with equality only for *i* = *j*. The corresponding metric distance is 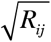. The main text uses the shorter phrase “effective distance” for *R*_*ij*_ ; geometrically it is the squared effective distance.

### C.3 The inner product in terms of effective distances

With a general reference residue *k*, define 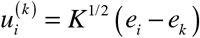. Direct expansion gives

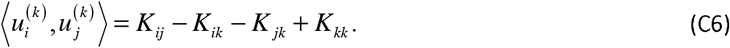

Now write each covariance difference in terms of *R*_*ab*_ = *K*_*aa*_ + *K*_*bb*_ − 2*K*_*ab*_. The diagonal terms cancel, leaving

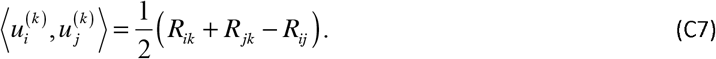

This is the discrete law of cosines. Quantities built only from the *R*_*ij*_ are unchanged under the Green-function replacement *G* = *K* + *c*1^*T*^ + 1*c*^*T*^. By contrast, *h* (*i*) − *h* (*j*) changes by 2(*c*_*i*_ − *c*_*j*_). The directional field in the main text is therefore tied to the Moore–Penrose convention, which is also the convention that identifies *h* (*i*) with the equilibrium mean-square displacement.

### C.4 The cooperation index

Take residue *i* as a reference and consider the relative displacement directions toward residues *j* and *k*. Their squared normalized overlap is

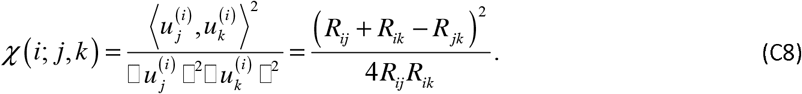

The second form follows by substituting Eq. (C7) with reference *i*, together with 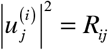 and 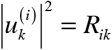. Equivalently, define the relative covariance

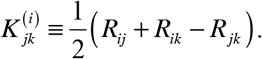

Then

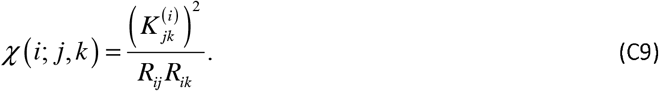

The ratio is dimensionless and invariant under a uniform rescaling of the network weights. By the Cauchy–Schwarz inequality,

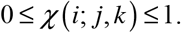

When χ is near 1, the two relative displacement directions are nearly aligned in the fluctuation geometry, consistent with strong sharing of structural constraints between the two channels. When χ is near 0, the two directions are nearly orthogonal, consistent with weak overlap between the two channels.

The same Gram construction extends to higher order. For three relative displacement vectors from a common reference, the Gram determinant is the squared volume of the parallelotope they span and provides a natural four-residue measure of collective geometry.

### C.5 Where order-three invariants enter

The specific triad invariant χ (*i*; *j, k*) does not enter the ordinary two-residue transfer-entropy expansion through the orders resolved in Appendix B. The leading asymmetry, Eq. (B15), closes on *h* and *S*_*ij*_. The equipotential correction, Eq. (B23), closes on *h, R*_*ij*_, *S*_*ij*_, and *L*_*ii*_ − *L*_*jj*_.

A three-residue invariant arises naturally when a third residue is included explicitly in the conditioned information measure. For example,

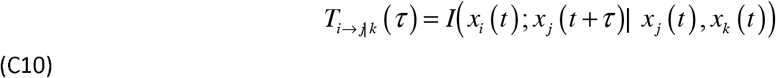

requires inversion of the present-state covariance block of (*i, j, k*). The denominator of that inverse is the determinant of the 3× 3 covariance block,

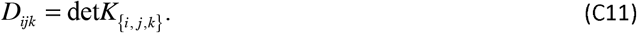

Thus triad structure belongs naturally to triad-conditioned transfer. It should not be inserted into the *O* (*τ* ^2^) pairwise result by analogy.

## Appendix D. Sign Persistence: Assumptions and Consequences

Appendix B fixes the initial direction of the transfer-entropy asymmetry as *τ* →0^+^. Whether that direction persists at finite delay is a separate question. Here sign persistence is stated precisely, and its consequences for finite-delay, integrated, and peak transfer are given directly.

### D.1 Definitions and assumptions

Assumption D.1 (Regularity). For each residue pair (*i, j*), the functions *T*_*i*→*j*_ (*τ*) and *T*_*j*→*i*_ (*τ*) are continuous on the compact interval [0,*τ*_max_]. Hence Δ*T*_*i*→ *j*_ (*τ*) is continuous and integrable and attains its extrema on that interval.

Assumption D.2 (Sign persistence). A nontrivial pair (*i, j*) is sign-persistent on [0,*τ*_max_] if there exists *σ*_*ij*_ ∈{−1, +1} such that

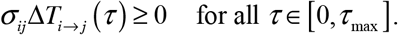

Equivalently,

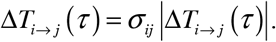

If Δ*T*_*i*→ *j*_ ≡ 0, the pair has no resolved directional asymmetry and is excluded from the statements below. In numerical work, sign persistence is tested only at the sampled delays. Reported persistence fractions therefore refer to the sampled relaxation window.

### D.2 The integral preserves the direction

Define the cumulative signed transfer

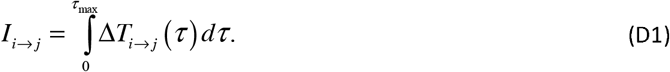

In the numerical work the integral is evaluated by the trapezoidal rule on the sampled grid, *τ*_min_ = 0.05 to *τ*_max_ = 5.0 ; the omitted interval 0 ≤ *τ* < *τ*_min_ contributes at order 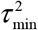 and does not affect any reported sign.

#### Proposition D.1.

If the pair is sign-persistent and

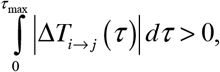

then sign*I*_*i*→ *j*_ = *σ*_*ij*_.

Proof. Sign persistence gives Δ*T*_*i*→ *j*_ (*τ*) = *σ*_*ij*_ Δ*T*_*i*→ *j*_ (*τ*). Because *σ*_*ij*_ is constant over the interval, it factors out of Eq. (D1). The remaining integral is positive by assumption. Hence the integral has sign *σ*_*ij*_ .⍰

Thus integration cannot reverse the direction of a sign-persistent asymmetry.

### D.3 Connection with the short-time expansion

Suppose

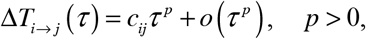

for sufficiently small positive *τ*. If *c*_*ij*_ ≠0 and the pair is sign-persistent near *τ* = 0, divide by the positive quantity *τ* ^*p*^ and take the right-sided limit. The sign of the finite-delay curve must then agree with the sign of *c*_*ij*_.

For the generic branch, Eq. (B15) gives *p* = 1 and

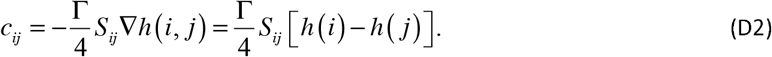

For *S*_*ij*_ > 0 and *h* (*i*) ≠*h* (*j*), sign persistence therefore implies

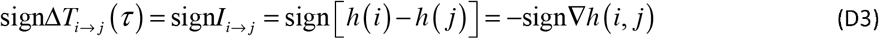

at every nonzero point of the sign-persistent window. If *S*_*ij*_ < 0, the leading sign is reversed. If *S*_*ij*_ = 0, the linear coefficient vanishes and Eq. (B15) does not define a leading direction.

For the equipotential branch, *h* (*i*) = *h* (*j*), Eq. (B23) gives *p* = 2 and

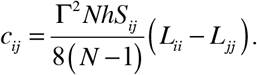

Thus, for *S*_*ij*_ > 0 and *L*_*ii*_ ≠*L*_*jj*_, a sign-persistent quadratic branch has the direction of *L*_*ii*_ − *L*_*jj*_.

### D.4 Peak transfer cannot reverse the direction

Let *F* (*τ*) = *T*_*i*→*j*_ (*τ*) and *G* (*τ*) = *T*_*j*→*i*_ (*τ*).

#### Proposition D.2.

If Δ*T*_*i*→*j*_ (*τ*) ≥0 throughout the window, then *F* (*τ*) ≥*G* (*τ*) pointwise and therefore

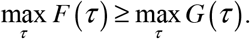

If Δ*T*_*i*→*j*_ (*τ*) ≤0, the inequality is reversed.

Proof. Suppose first that *F* (*τ*) ≥*G* (*τ*) for all *τ*. By continuity on a compact interval, *G* attains its maximum at some *τ*_0_. At that same point, *F* (*τ*_0_) ≥*G* (*τ*_0_) = max_*τ*_ *G* (*τ*), while max_*τ*_ *F* (*τ*) ≥*F* (*τ*_0_). Combining the two inequalities gives the result. The opposite-sign case follows by interchanging *F* and *G* .⍰

Hence sign persistence prevents the peak comparison from indicating the opposite direction. Exact equality remains possible; a strict peak inequality requires an additional nondegeneracy condition.

### D.5 Positive normalizations preserve direction

Let *A*_*i*_ > 0 and *A*_*j*_ > 0 be positive normalizing factors. Since division by a positive quantity does not change a sign,

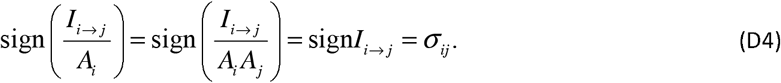

In particular, for the leading branch with *S*_*ij*_ > 0 and *h* (*i*) ≠*h* (*j*), divide Eq. (B15) by its positive magnitude factor Γ*τ S*_*ij*_ /4 and choose the opposite orientation so that the normalized field points along the potential gradient:

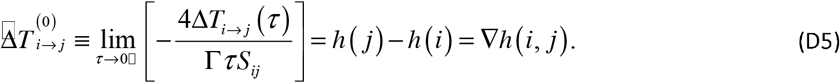

The minus sign is therefore an orientation convention, not an additional dynamical assumption. The normalization applies only to the linear branch. For equipotential pairs the linear normalization is indeterminate, and the leading direction is given by Eq. (B24).

### D.6 Scope and empirical status

Sign persistence is not a theorem of the Gaussian model and is not a thermodynamic entropy current. It is a property of a residue pair over a specified delay interval and must be checked from the finite-delay transfer curves. When it holds, the finite-delay asymmetry, its integral, and the peak comparison are directionally consistent with the leading short-time term.

The KRAS calculation shows that persistence is common but not universal. Evaluating Δ*T*_*i*→*j*_ (*τ*) at 100 equally spaced lags from *τ* = 0.05 to *τ* = 5.0, 96.86% of the 796 contacts retain their short-time sign. Over all 14,706 residue pairs, the short-time sign agrees with the sign of the integrated asymmetry in 98.81% of cases. These percentages refer to different pair sets and measure different properties.

The Gln61 → Gly60 contact retains its direction over the sampled window, as do the leading contacts generally. The 25 contacts that reverse are, without exception, weak. The largest, Lys128 → Thr127, carries an integrated asymmetry of 7.75×10^−4^ nats, two orders of magnitude below the strongest contacts; the set also includes the large-separation pairs Glu49 → Ser1 and Lys5 → Glu76. Reversal is therefore confined to channels too weak to be resolved against the leading structure of the map.

The short-time direction should therefore not be extrapolated to long delay without checking the exact transfer curves.

