## Supplementary Information for "Flexibility Drives Information Flow in Proteins: Fluctuation Potential Gradients Dictate Directional Entropy Transfer"

The notation is that of the main text and of Appendices A–D. Each quantity refers to one Cartesian component of the displacement, and entropies are expressed in nats. Nothing enters the calculation but the deposited coordinates.

### S1. Model, parameters and numerical checks

The chain is represented by its 172 Cα positions. Two residues are joined whenever they lie within $r_c$ of one another, and the junction is assigned the weight $\gamma_{ij} = \exp(-d_{ij} / T_W)$. Close neighbours are thereby stiffly coupled and distant ones only loosely, in contrast to the uniform coupling of the elementary network. The weighted Kirchhoff matrix L follows at once from these weights, and its pseudoinverse K is the covariance of the fluctuations. Table S1 collects the constants of the model and the quantities derived from them.

Table S1. Model parameters, derived quantities and numerical checks.

| **Quantity** | **Value** |
|---|---|
| Structure, chain | 6GOD, A |
| Residues, N | 172 |
| Contact cutoff, $r_c$ | 7.8 Å |
| Weighting scale, $T_W$ | 1.0 Å |
| Weighted contacts | 796 |
| Distinct residue pairs | 14,706 |
| Relaxation-rate prefactor, Γ | 1 |
| Short-time delay, $\tau_{short}$ | 0.01 |
| Lag window | $0.05 \le \tau \le 5.0$ |
| Lag points | 100, equally spaced |
| $\lambda_{max}$ | 0.1293 |
| $\tau_{short} \lambda_{max}$ | $1.293 \times 10^{-3}$ |
| max $\lvert LK - P \rvert$ | $9.6 \times 10^{-15}$ |
| min $\lvert \rho_{ij} \rvert$ over contacts | $1.048 \times 10^{-2}$ |
| Threshold $1/(N - 1)$ | $5.848 \times 10^{-3}$ |
| Contacts with $S_{ij} > 0$ | 796 of 796 |
| Sign-persistent contacts | 771 of 796 (96.86 %) |
| sign(I) = short-time sign, all pairs | 98.81 % |
| Spearman $r_s$, h vs $B_{exp}$ | 0.807 |
| Pearson r, h vs $B_{exp}$ | 0.750 |

| | |
|---|---|
| Contacts agreeing with ∇B_exp | 77.64 % |

### S2. Validity of the short-time expansion

The expansion of the matrix exponential is legitimate so long as $\Gamma\tau\lambda\_max$ remains small compared with unity. This product is $1.29 \times 10^{-3}$ at $\tau = 0.01$, and the linear term of Eq. (11) accordingly governs the map of Figure S1. At the longer delays of Sections S3, S4 and S7 no expansion is made. The transfer entropy is evaluated there from the Schur complements of $K(\tau) = \exp(-\Gamma L\tau)K$, which is exact for a Gaussian network at every delay.

The two routes agree as they must. Compared at $\tau = 10^{-4}$, $10^{-3}$ and $10^{-2}$, the exact result departs from Eq. (B15) by an amount proportional to τ, which is the behaviour required of a neglected term of second order. Eq. (B13) behaves likewise.

### S3. Strongest directional contacts

The asymmetry of each of the 796 junctions, integrated over the relaxation window, measures the net communication carried by that junction; its sign names the source. The ten largest are given in Table S2. Gln61 → Gly60 stands first.

Table S2. The ten strongest integrated directional contacts.

| Source → sink | \|i − j\| | I (nats) | S_ij | R_ij | h(src) | h(sink) |
|---|---|---|---|---|---|---|
| Gln61 → Gly60 | 1 | $1.841 \times 10^{-2}$ | $4.49 \times 10^{-4}$ | 29.90 | 43.06 | 27.61 |
| His166 → Ile163 | 3 | $1.776 \times 10^{-2}$ | $2.98 \times 10^{-4}$ | 36.36 | 62.99 | 42.85 |
| Val109 → Pro110 | 1 | $1.775 \times 10^{-2}$ | $3.65 \times 10^{-4}$ | 32.91 | 43.94 | 27.37 |
| Lys165 → Ile163 | 2 | $1.766 \times 10^{-2}$ | $3.82 \times 10^{-4}$ | 31.19 | 58.50 | 42.85 |
| Lys167 → Ile163 | 4 | $1.764 \times 10^{-2}$ | $2.01 \times 10^{-4}$ | 46.20 | 73.01 | 42.85 |
| Arg164 → Ile163 | 1 | $1.751 \times 10^{-2}$ | $5.87 \times 10^{-4}$ | 23.72 | 53.24 | 42.85 |
| Lys169 → His166 | 3 | $1.714 \times 10^{-2}$ | $2.43 \times 10^{-4}$ | 37.98 | 87.42 | 62.99 |
| Thr74 → Gly75 | 1 | $1.702 \times 10^{-2}$ | $4.98 \times 10^{-4}$ | 27.83 | 41.57 | 29.70 |
| Lys147 → Ala146 | 1 | $1.684 \times 10^{-2}$ | $5.85 \times 10^{-4}$ | 25.48 | 40.00 | 29.78 |
| Met170 → His166 | 4 | $1.658 \times 10^{-2}$ | $1.70 \times 10^{-4}$ | 48.27 | 97.56 | 62.99 |

In every instance the source is the residue of larger node potential, as Eq. (11) requires, and in every instance the direction so assigned is maintained throughout the window.

### S4. Residue-level sources and sinks

Summed over the junctions that meet at a residue, the integrated asymmetry gives the net flux $F_i$ borne by that residue. A positive value marks a residue that supplies information to its neighbours, a negative one a residue that receives it.

Table S3. The five leading sources and the five leading sinks, with rank out of 172.

| Sources | Rank | F_i (nats) | Sinks | Rank | F_i (nats) |
|---|---|---|---|---|---|
| Ser1 | 1 | $+4.77 \times 10^{-2}$ | Ser145 | 172 | $-7.67 \times 10^{-2}$ |
| Thr148 | 2 | $+4.49 \times 10^{-2}$ | Gln22 | 170 | $-5.11 \times 10^{-2}$ |
| Gln25 | 3 | $+4.27 \times 10^{-2}$ | Ile163 | 171 | $-5.17 \times 10^{-2}$ |
| Lys172 | 4 | $+4.10 \times 10^{-2}$ | Leu52 | 167 | $-4.73 \times 10^{-2}$ |
| Gly48 | 6 | $+3.89 \times 10^{-2}$ | Ala134 | 163 | $-4.37 \times 10^{-2}$ |

Ser1, Lys172 and Ser171 are ends of the chain. An end is unconstrained on one side for a trivial reason, and its large potential reflects that circumstance rather than the architecture of the fold; the main text therefore quotes the leading interior residues. Gly60 stands 162nd and Phe156 53rd.

### S5. Sequence separation

Divided at $|i - j| = 3$, the junctions fall into 424 local and 372 nonlocal. The local ones carry the larger asymmetries, 78 of the first hundred being of this kind. Nothing else is to be expected. Neighbours along the chain share the largest covariance and are separated by the smallest effective distance, and the coupling factor S_ij is correspondingly large.

It is the nonlocal set that reports on the fold. Its leading members are the i, i+4 contacts of the C-terminal helix — Lys167 → Ile163, Met170 → His166, Lys169 → Lys165 — and, at true tertiary separation, Val45 → Cys51, Ile46 → Cys51 and Ser1 → Leu52. The two contacts on Cys51 constitute the passage from Switch I to Switch II described in Section 3.3.

### S6. Controls

It must be asked whether the agreement with the observed B-factors expresses the architecture of the molecule or merely the number of contacts each residue happens to make. Three comparisons answer this. In the first, every weight is set to unity, so that only the pattern of contacts survives. In the second, h(i) is compared directly with the number of contacts. In the third, the contacts are rearranged at random with the degree of every residue held fixed and the original weights redistributed among them, a thousand times over.

Table S4. Controls on the B-factor agreement.

| **Model** | **Statistic** | **Value** |
|---|---:|---:|
| Weighted network | Spearman r_s, h vs B_exp | 0.807 |
| Weighted network | contacts agreeing with ∇B_exp | 77.64 % |
| Uniform conductance | Spearman r_s, h vs B_exp | 0.815 |
| Contact degree | Spearman r_s, h vs degree | −0.786 |
| Rewired null, 1000 replicates | mean agreement | 66.08 % |
| Rewired null, 1000 replicates | standard deviation | 2.25 % |
| Rewired null, 1000 replicates | largest replicate | 73.24 % |
| Rewired null, 1000 replicates | z-score of observed value | 5.15 |

Not one of the thousand rearrangements attains the observed 77.64 per cent. The weighted and the uniform networks rank the residues almost identically against B_exp; the weighting alters the magnitudes of the potentials but not their order, which is fixed by the pattern of contacts. That h(i) falls as the number of contacts rises is the elementary mechanical statement that a residue held on many sides moves least.

### S7. Lag dependence and sign persistence

Two figures are quoted in the main text and they are not the same figure. Of the 796 junctions, 771, or 96.86 per cent, hold the sign given them at short times at every delay sampled between 0.05 and 5.0. Of all 14,706 pairs, 98.81 per cent have an integrated asymmetry whose sign agrees with the short-time one. The first statement concerns the whole curve, the second only its integral, and neither follows from the other.

The delay dependence of the three strongest junctions is given in Table S5. Each rises steadily across the window, so that the direction fixed at short times accumulates without cancellation.

Table S5. Exact ΔT_i→j at selected delays (nats).

| τ | Gln61 → Gly60 | His166 → Ile163 | Val109 → Pro110 |
|---|---|---|---|
| 0.05 | $8.658 \times 10^{-5}$ | $7.493 \times 10^{-5}$ | $7.562 \times 10^{-5}$ |
| 0.10 | $1.728 \times 10^{-4}$ | $1.498 \times 10^{-4}$ | $1.511 \times 10^{-4}$ |
| 5.00 | $6.709 \times 10^{-3}$ | $6.898 \times 10^{-3}$ | $6.823 \times 10^{-3}$ |

The twenty-five junctions that reverse, 3.14 per cent of the whole, are without exception weak ones. The largest of them, Lys128 → Thr127, carries $7.75 \times 10^{-4}$ nats, smaller by two orders of magnitude than the junctions of Table S2, and several are pairs of large sequence separation such as Glu49 → Ser1 and Lys5 → Glu76. Reversal is thus confined to channels too feeble to be read, and the map is unaffected at the magnitudes that are interpreted.

### S8. Supplementary figures

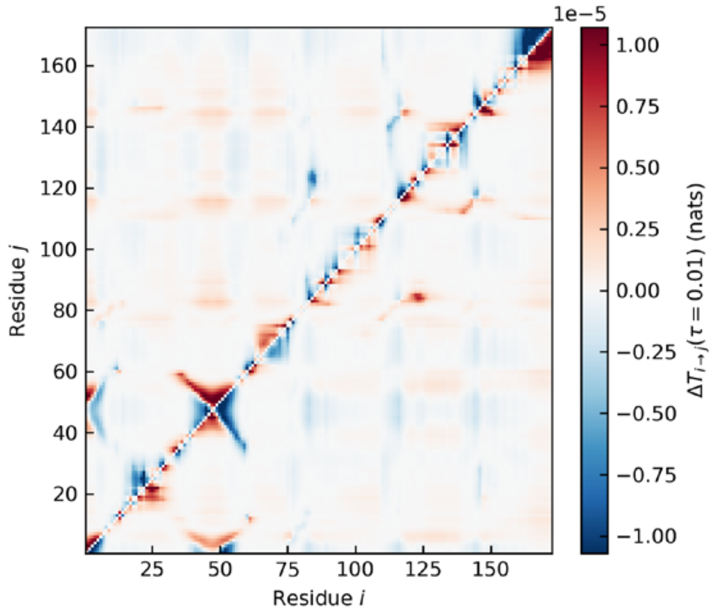


Figure S1. Short-time directional asymmetry ΔT_i→j at τ = 0.01 for all residue pairs, without thresholding. Colour limits are the 99.5th percentile of |ΔT|. Figure 1 of the main text is the quartile-thresholded integrated version.

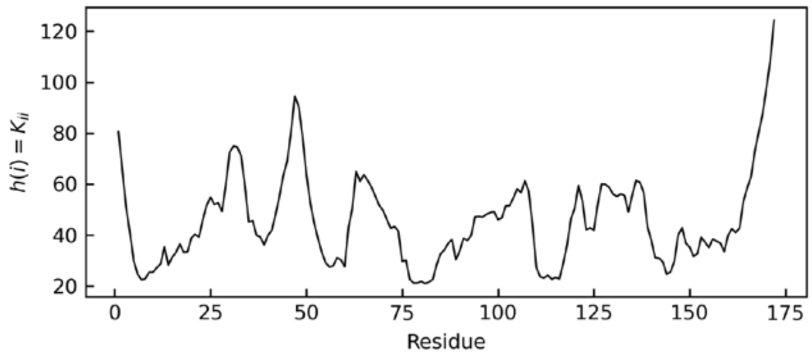

Figure S2. Residue fluctuation potential h(i) = K_ii along the chain. The maxima lie at the termini and the flexible loops.

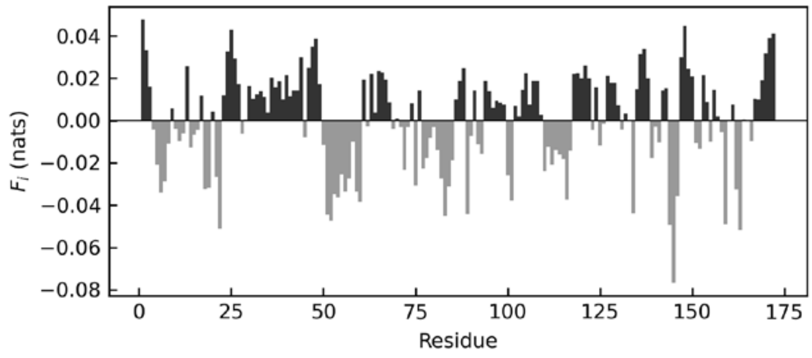


Figure S3. Net integrated flux F_i summed over the 796 weighted contacts. Positive bars are sources, negative bars sinks.

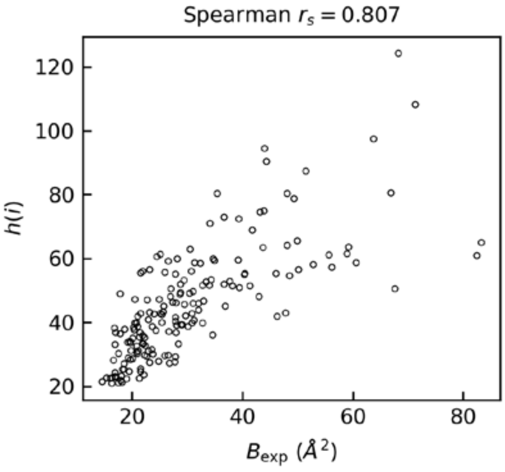


Figure S4. Calculated h(i) against the experimental Cα B-factor of 6GOD. Spearman r_s = 0.807.

### S9. Reproduction script and machine-readable files

A single script, entropy_transfer_6GOD.py, reproduces every number, table and figure above from the coordinate entry alone. No parameter is fitted and no trajectory is required.

Tables. TableS1_parameters.csv (parameters and checks), TableS2_strongest_contacts.csv (all 796 contacts with I, the short-time asymmetry, S_ij, R_ij, both node potentials and the persistence flag), TableS3_residues.csv (all 172 residues with h, B_exp, F_i, contact degree and the uniform-weight potential), TableS4_lag_dependence.csv (100 delays for three representative contacts), all_pairs_report.csv (all 14,706 pairs with K_ij, R_ij, S_ij, ρ_ij and both asymmetries), null_replicates.csv (1000 rewired replicates).

Figures. Figure1_integrated_map.tif is Figure 1 of the main text. FigureS1_short_time_map.tif, FigureS2_node_potential.tif, FigureS3_flux.tif and FigureS4_bfactor.tif are the four supplementary figures of Section S8.
